# Human patient *SFPQ* homozygous mutation is found deleterious for brain and motor development in a zebrafish model

**DOI:** 10.1101/2020.03.18.993634

**Authors:** Stephanie Efthymiou, Patricia M. Gordon, Vincenzo Salpietro, Triona Fielding, Eugenia Borgione, Carmela Scuderi, Henry Houlden, Corinne Houart

## Abstract

SFPQ (Splicing factor proline- and glutamine-rich) is a DNA and RNA binding protein involved in transcription, pre-mRNA splicing, and DNA damage repair and it has been previously implicated in neurodegenerative disorders. A homozygous p.Ser660Asn variant in *SFPQ* was identified through whole exome sequencing (WES) in an Italian woman presented a complex neurological phenotype with intellectual disability, peripheral neuropathy, bradykinesia, extrapyramidal rigidity and rest (heads) tremor and neuroradiological anomalies including thin dysplastic corpus callosum, hypomyelination and hypointensity of the globus pallidus and of the mesencephalic substantia nigra (resembling neurodegeneration with brain iron accumulation; NBIA). Using a zebrafish *SFPQ* genetic model we have showed that a rescue with this SFPQ^S660N^ mutant revealed robust defects in the developing central nervous system (CNS) of the embryos, including abnormal branching of the motor axons innervating body muscles and misfolding of the posterior brain neuroepithelium. The defects hereby identified in the model organism indicate a potential contribution of the homozygous *SFPQ* p.Ser660Asn variant in some of the patient’s neurodegenerative features, including the clinical parkinsonism and the *NBIA-like* pattern on brain imaging.

## Introduction

The development of the human brain relies on complex mechanisms involving both the generation of appropriate cell types and their organization in the correct anatomical structures. Post-transcriptional regulation of gene expression plays a fundamental role in the temporal and spatial modulation of early development and has been recently found to be involved in many disease mechanisms.

Cerebellar dysfunction is a recurrent feature of several neurodevelopmental disorders, including intellectual disability (ID), autism spectrum disorders (ASDs), developmental epileptic encephalopathies (DEEs) and speech disturbances and abnormal cerebellum development (during early embryonic stages) can significantly contribute in the anomalies of movement, cognition, and affective regulation frequently observed within the phenotypic spectrum of these disorders (Stoodley, 2016). Biallelic mutations in genes implicated in RNA processing and metabolism can affect critical post-transcriptional events since the earliest stages of life (Nachtergaele and He, 2017). Understanding the impact of mutations in such genes on neurodevelopment and disease phenotypes will allow us to elucidate initiating molecular events as well as guide the development of new therapies targeting primary mechanisms before the disease progresses too far.

*SFPQ* encodes for a Splicing Factor Proline/Glutamine-Rich (Sfpq) multifunctional protein, ubiquitously expressed across the body from embryo to adult in all vertebrates and known to be involved in transcriptional regulation, pre-mRNA splicing and RNA transport (Bottini et al., 2017; Cosker et al., 2016; Shav-Tal and Zipori, 2002; Takeuchi et al., 2018). Human *SFPQ* lies in a region on chromosome 1p34-p36 already implicated in speech anomalies and language impairment and has been found to be mis-regulated in brains of patients with a variety of neurodegenerative disorders such as autism and dyslexia (Chang et al., 2015; Ke et al., 2012; Stamova et al., 2013; Tapia-Paez et al., 2008). The SFPQ protein is conserved across species and plays a key role in neuronal development and network organization (Thomas-Jinu et al., 2017). The protein structure contains tandem RNA recognition motif domains, a NOPS domain, a coiled-coil region and an N-terminal proline/glutamine-rich low-complexity region (Passon et al., 2012).. More recently, loss of SFPQ function has been implicated as a risk factor for human neurodegenerative diseases such as amyotrophic lateral sclerosis (ALS), fronto-temporal dementia (FTD) and Alzheimer’s Disease (AD) (Ishigaki et al., 2017; Luisier et al., 2018; Takayama et al., 2019; Thomas-Jinu et al., 2017) in mouse, iPSC, and zebrafish models. Antibody staining in zebrafish showed that the protein is localised in nuclei in all tissues throughout development and in neurites of a subset of neuronal population, including motor neurons. This unique cytoplasmic pool was found to drive motor neuron maturation in absence of nuclear function (Thomas-Jinu et al., 2017).

Recent RNA sequencing studies of patient iPSC-derived neuron cultures from both familial and sporadic ALS patients showed abnormal premature intron retention in the *SFPQ* transcript during early neural differentiation across the mutant cultures, leading to loss of function of SFPQ. These results revealed SFPQ loss of function as an early hallmark of ALS (Luisier et al., 2018). Here, we report the finding of a link between a severe genetic neurodevelopmental disorder and a homozygous missense mutation in *SFPQ*. We investigated the exome of a female patient presented with microcephaly, neuropathy, head tremor and severe cerebellar atrophy with nystagmus and dysarthria. The patient is a 47-year-old woman, born at term by normal delivery from healthy Sicilian parents non-consanguineous for their account (Figure 1A-B). The patient’s family history was positive for intellectual disability in her first-degree cousin. Her birth weight, length and Apgar scores have not been reported. Since the first months of her life, the patient presented muscle hypertonia and a severe delay in psychomotor development with poor spontaneous movements, lack of head control, frontal plagiocephaly, weak crying, and absent suction. An episode of seizures triggered by fever was reported at the age of 1 year. One year later, she presented an episode of (seizure-free) generalized tonic-clonic seizures. Since early adulthood she developed recurrent partial epileptic episodes and a treatment with valproic acid was started at time, with partial control of seizures. These episodes were frequently accompanied by ocular staring and brief loss of consciousness. The patient was first admitted to the IRCCS Oasi Maria SS. Troina Medical center (Troina, Italy) at the age of 38 years old. At this time, physical examination revealed microcephaly, edentulism, globose abdomen, and skin trophic alterations. Neurological examination revealed spastic tetraparesis and ataxia with ocular motor apraxia, also featured by marked deficits of horizontal eye movements and nystagmus. At this time, she also showed signs of bradykinesia, extrapyramidal rigidity (with abnormally increased resistance to movement) and rest tremor, mainly involving the head (See supplementary Video 1). Babinski reflex was absent and also deep tendon reflexes in the four limbs were not elicitable. Dysmetria and intention tremor were present in the upper limbs. Muscle wasting was also predominant in her lower limbs with retraction of hip adductor group and triceps surae muscles. A neuropsychological assessment as part of her neurological evaluation revealed a severe intellectual disability. Electromyography (EMG) showed motor and sensory axonal polyneuropathy. Electroencephalogram (EEG) highlighted focal diffuse paroxysmal activity over the frontal and the parietal regions. Brain magnetic resonance imaging (MRI) performed at the age of 40 years old showed severe cerebellar atrophy, associated to some brainstem atrophy, and thin dysplastic corpus callosum (Figure 1E). In the supratentorial area, white matter atrophy and signal changes were also documented. A symmetric hypointensity of the globus pallidus and of the mesencephalic substantia nigra was also documented, suggesting anomalous deposit of paramagnetic substances (e.g., iron). Brain computed tomography (CT) scan did not showed intracranial calcifications. Ophthalmological and fundoscopy examinations identified bilateral optic atrophy and strabismus. Visual evoked potentials (VEPs) appeared prolonged. Electrocardiogram (ECG) and echocardiogram were normal. The abdominal ultrasound described an unspecified hepatopathy. Urine organic acids, karyotype and array comparative genome hybridization were all normal. Panel NGS analysis for 12 genes implicated in brain neurodegeneration and iron accumulation did not show any pathogenic mutations or intragenic deletions/duplications. Muscle biopsy suggested a picture suggestive of vacuolar myopathy associated to areas of adipose fat replacement, neurogenic dysfunction, and denervation. Also, a reduction of respiratory chain complex I and II was documented. At the follow-up appointments, a progression of the neuromotor symptoms, featuring an increased rigidity to the passive movements and increase of the tremor, has been observed. Follow-up brain imaging studies confirmed the previous radiological findings (not shown).

**Figure 1.**
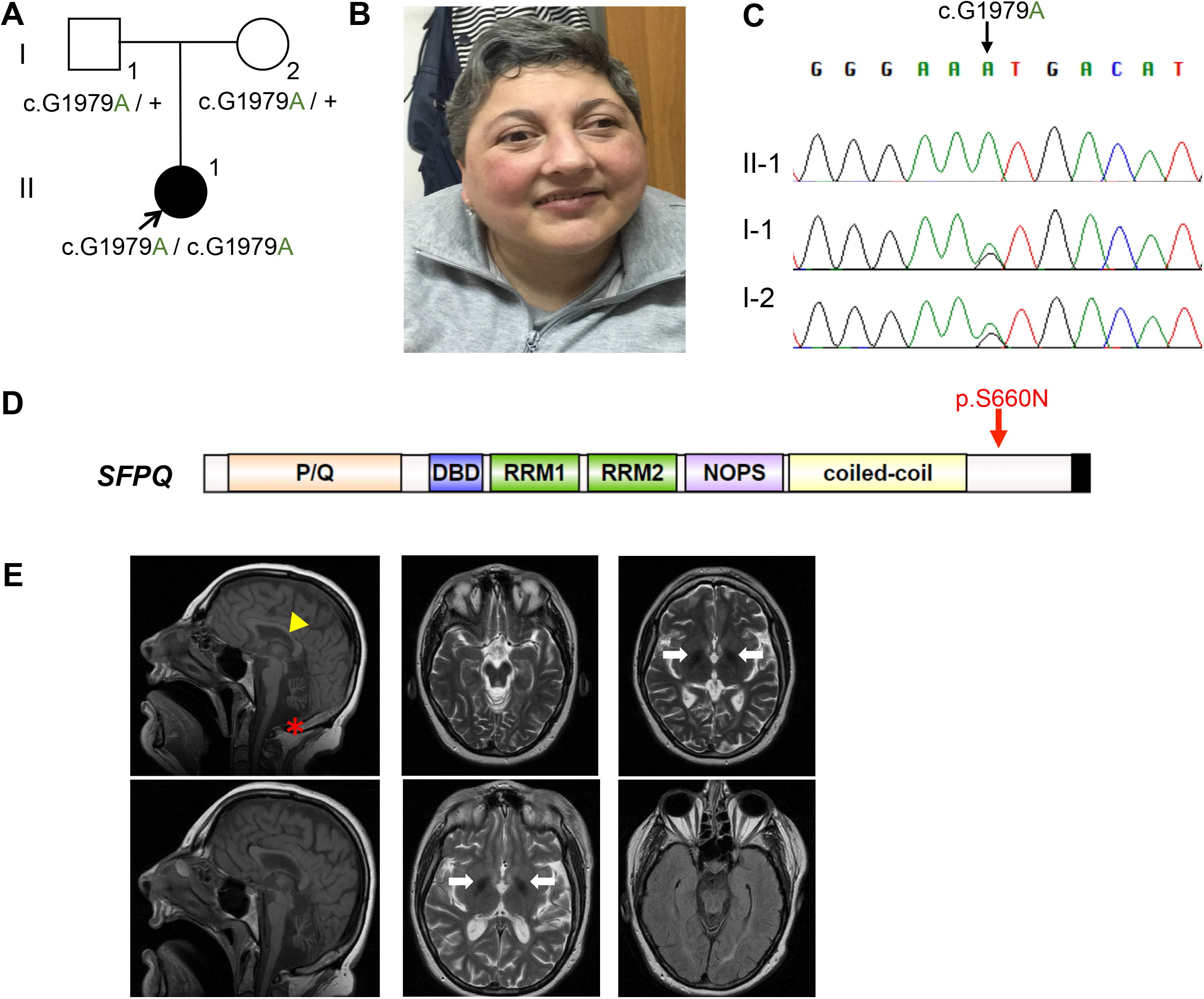
**A.** Pedigree of the Italian family showing the inheritance of the *SFPQ* variant in proband and parents. **B.** Facial photograph of the 47-year-old patient, presented with lack of head control, frontal plagiocephaly, head tremor, ocular motor apraxia and marked deficits of horizontal eye movements and nystagmus. **C.** Segregation analysis of the *SFPQ* variant by traditional Sanger sequencing, showing the *SFPQ* variant identified as homozygous in the proband and heterozygous in the two parents. **D.** Protein structure of SFPQ, made up of important domains mentioned above, and showing the location of the *SFPQ* variant identified. **E.** Top panel: MRI studies at the age of 36 years. Sagittal T1 and axial T2 studies reveal revere cerebellar and vermian atrophy, with brainstem ipotrophy (red asterisk) and thinning of the corpus callosum (yellow triangle). In the supratentorial area, periventricular white matter signal abnormalities are evident. Symmetric hypointensity of the globus pallidus is also present (white arrows). Bottom panel: At the age of 40 years abnormal white matter signal, cerebellar and vermian atrophy and brainstem hypotrophy is documented in T1 axial images. Hypointensity of globus pallids in T2 and inversion recovery fast spin◻echo flair sequences is also detected.

## Materials and Methods

### Whole exome sequencing (WES)

After informed consent, blood samples were collected from the patient and her parents, and extracted DNA using standard procedures. To investigate the genetic cause of the disease, WES was performed in both the affected female and the two parents (Figure 1A: II-1, I-1, I-2). Nextera Rapid Capture Enrichment kit (Illumina) was used according to the manufacturer instructions. Libraries were sequenced in an Illumina HiSeq3000 using a 100-bp paired-end reads protocol. Sequence alignment to the human reference genome (UCSC hg19), and variants calling, and annotation were performed as described elsewhere (Mencacci et al., 2016). In total, 55,531,100 (II-1) unique reads were generated. After removing all synonymous changes, we filtered single nucleotide variants (SNVs) and indels, only considering exonic and donor/acceptor splicing variants. In accordance with the pedigree and phenotype, priority was given to rare variants [<1% in public databases, including 1000 Genomes project, NHLBI Exome Variant Server, Complete Genomics 69, and Exome Aggregation Consortium (ExAC v0.2)] that were fitting a recessive (e.g., compound heterozygous or homozygous) or a *de novo* model.

### Zebrafish handling and maintenance

Zebrafish (*Danio rerio*) were reared in accordance with the Animals (Scientific Procedures) Act 1986. Fish were maintained at 28C, and embryos were cultured in fish water containing 0.01% methylene blue to prevent fungal growth. Wildtype fish were from the AB strain.

### RNA injections

To make RNA transcripts for hSFPQ, hSFPQ^S660N^, gfp-hSFPQ, and gfp-hSFPQ^S660N^, DNA sequences were inserted into the plasmid pCS2+ (Addgene). Plasmids were linearized and transcribed using the mMessage mMachine SP6 Transcription Kit (Ambion). Capped RNA was purified using mini Quick spin columns (Roche). Embryos were injected at the one-cell stage with 150 pg of hSFPQ, hSFPQ^S660N^ RNA or 130 pg of gfp-hSFPQ, and gfp-hSFPQ^S660N^ RNA.

### Whole mount immunohistochemistry

Embryos were fixed in 4% paraformaldehyde overnight at 4°C, then washed with PBS+0.8% triton X-100, permeabilized with 0.25% trypsin/PBS, and blocked in 10% goat serum/PBS for one hour. Embryos were incubated in primary antibody at 4°C overnight, washed with PBS, and incubated with secondary antibody at 4°C overnight. Primary antibodies were anti-gfp 1:500() and anti-acetylated tubulin 1:1000 (). Secondary antibodies were Alexa Fluor 488/568 (Life Technologies). Alexa Fluor 568 phalloidin stain (Life Technologies) was added at 1:1000 during secondary antibody incubation.

### In-situ hybridization

*In-situ* hybridization was performed as previously described (Thomas-Jinu et al, 2017). Antisense probes targeting the genes *Otx2 and Gbx2* were created from linearized plasmids using an *in-vitro* transcription reaction according to the manufacturer’s instructions using DIG Labelling Mix (Roche) or fluorescein Labelling Mix (Roche). Probes were purified using mini Quick spin columns (Roche) and stored in 5X SSC and 50% formamide. Embryos were fixed overnight at 4°C in 4% PFA and washed in PBS/0.1% Tween-20 (PBST), then dehydrated in methanol and stored at −20°C. For hybridization, embryos were rehydrated in PBST, then treated with 10 ug/ml Proteinase K (Sigma) before further fixation in 4% PFA for 20 minutes. Embryos were then incubated for several hours at 65°C in hybridization buffer: 5xSSC, 1% Tween-20, 0.5 mg/ml torula RNA, 50μg/ml heparin, 0.1% CHAPS, 50% formamide. Embryos were then incubated overnight in probes diluted 1:20 in hybridization buffer. The following day, embryos were washed over several hours at 65°C in first hybridization buffer, then 2XSSC/1%CHAPS, then 0.2%SSC/1%CHAPS, then at room temperature in MAB/0.1%Tween. They were blocked in MAB/0.1%Tween + 2% Blocking Reagent (Roche). Embryos were then incubated overnight at 4°C with 1:4000 anti-digoxigenin (Roche) in blocking solution. After washing for several hours in MAB/0.1% Tween at room temperature, embryos were developed in NBT + BCIP in 0.1M NaCl, 0.1M Tris-Hcl pH 9.5, 0.05M MgCl2, 1% Tween-20. After development, embryos were fixed in 4% PFA for 75 minutes.

## Results

Whole-exome sequencing of the trio revealed a heterozygous de novo mutation (NM:004321: c.920G>A; p.Arg307Gln) affecting a conserved residue within the motor domain (MD) of the *KIF1A* gene. In addition, a homozygous missense variant in the *SFPQ* gene (NM_005066:c.G1979A; p.Ser660Asn) was also identified. The p.Ser660Asn *SFPQ* variant involves a conserved Serine residue and was present within the most significant homozygous block (chr1: 33549405-40840270) identified by homozygosity mapping analysis (which was performed using the WES data). The variant is absent from GnomAD (gnomad.broadinstitute.org) containing 125,748 exome sequences from unrelated individuals, and it is considered pathogenic by several in silico predictors (including Mutation Taster, FATHMM, Sift, LRT, Eigen, MetaLR). Both parents were found to be heterozygous carriers of the p.Ser660Asn variant in Sanger-based segregation analysis (Figures 1C-D). The p.Arg307Gln *KIF1A* variant could partially explain the complex neurological phenotype of our patient as established in ClinVar (www.ncbi.nlm.nih.gov/clinvar; accession number: 418275) and in literature reports as a disease-causing mutation, implicated in phenotypes ranging between intellectual disability, spasticity, optic nerve and/or cerebellar atrophy (Cherot et al., 2018; Hotchkiss et al., 2016; Ohba et al., 2015). Thus, the rare homozygous *SFPQ* variant emerged as a potential candidate genetic variant to explain part of the complex clinical neurological features not previously associated to the *KIF1A*-related neurodevelopmental disorder, such as the bradykinesia, rest (head) tremor and extrapyramidal rigidity, as well as the MRI findings suggestive of neurodegeneration with brain iron accumulation (NBIA). This is also supported by the predicted severe impact of the *SFPQ* p.Ser660Asn mutation on protein function. Previous reports implicating *SFPQ* missense variants as a genetic risk factor for ALS and neurodegenerative diseases (Thomas-Jinu et al., 2017) where frontotemporal lobar degeneration (FTLD) patients have developed brain iron accumulation in the basal ganglia (De Reuck et al., 2014).

To address whether the homozygous missense mutation found in our patient may be causative for her clinical parkinsonism and the “*NBIA-like”* features identified by repeated brain imaging studies, we used a zebrafish *SFPQ* genetic model. We previously showed that *sfpq* null loss of function homozygous mutation leads to brain boundary defects, cerebellar abnormalities and loss of motor function in zebrafish (Thomas-Jinu et al., 2017) and found a link between this gene and Amyotrophic Lateral Sclerosis (ALS). We have shown that human full-length human *SFPQ* is able to rescue completely the loss of gene function in zebrafish. We therefore tested whether the human SFPQ^S660N^ transcript had the ability to rescue the zebrafish mutant and if so, whether this rescue was accompanied with neurodevelopmental abnormalities. Rescue experiments are done blind by injection of full-length wild-type hSFPQ or hSFPQ^S660N^ transcripts at 1-cell stage (SFPQ is expressed ubiquitously from fish to human) into an incross of *sfpq*^+/−^ zebrafish at the one-cell stage (25% of the embryos are homozygous null), followed by pan-axonal staining at 36 or 48 hours post-fertilisation (hpf). We first performed a set of double-blind experiments to assess whether any visible abnormalities were associated with rescue of the null mutant by the S660N allele (Suppl. Figure 1). Homozygous mutant embryos rescued with SFPQ^S660N^ showed robust abnormalities in motor innervation (Suppl. Figure 1B), so we further investigated the developing CNS in these embryos using phalloidin stain and anti-acetylated tubulin to visualize the cytoskeleton (Figure 2). This analysis confirmed abnormal branching of the motor axons innervating body muscles (Figure. 2J-L,N,O) and a spectacular misfolding of the neuroepithelium at the isthmus, bulging either into the tectal area (equivalent to superior colliculus) or into the rostral brain stem (Figure 2 A-L) in addition to reduction of telencephalic volume (Figure 2P) and reduction of the distance between eyes (Figure. 2Q), although the eyes themselves are normal (Figure 2M). The isthmic dysmorphology is accompanied by a lack of differentiation of the cerebellum. To identify whether the misfolded tissue was of midbrain or hindbrain identity, we performed *Otx2* (midbrain marker), *Gbx2* (hindbrain marker) double *in situ* mRNA hybridization and found that the ectopic lump was of dual origin, comprising *Otx2-* and *Gbx2*-expressing cells (Figure 3).

**Figure 2.**
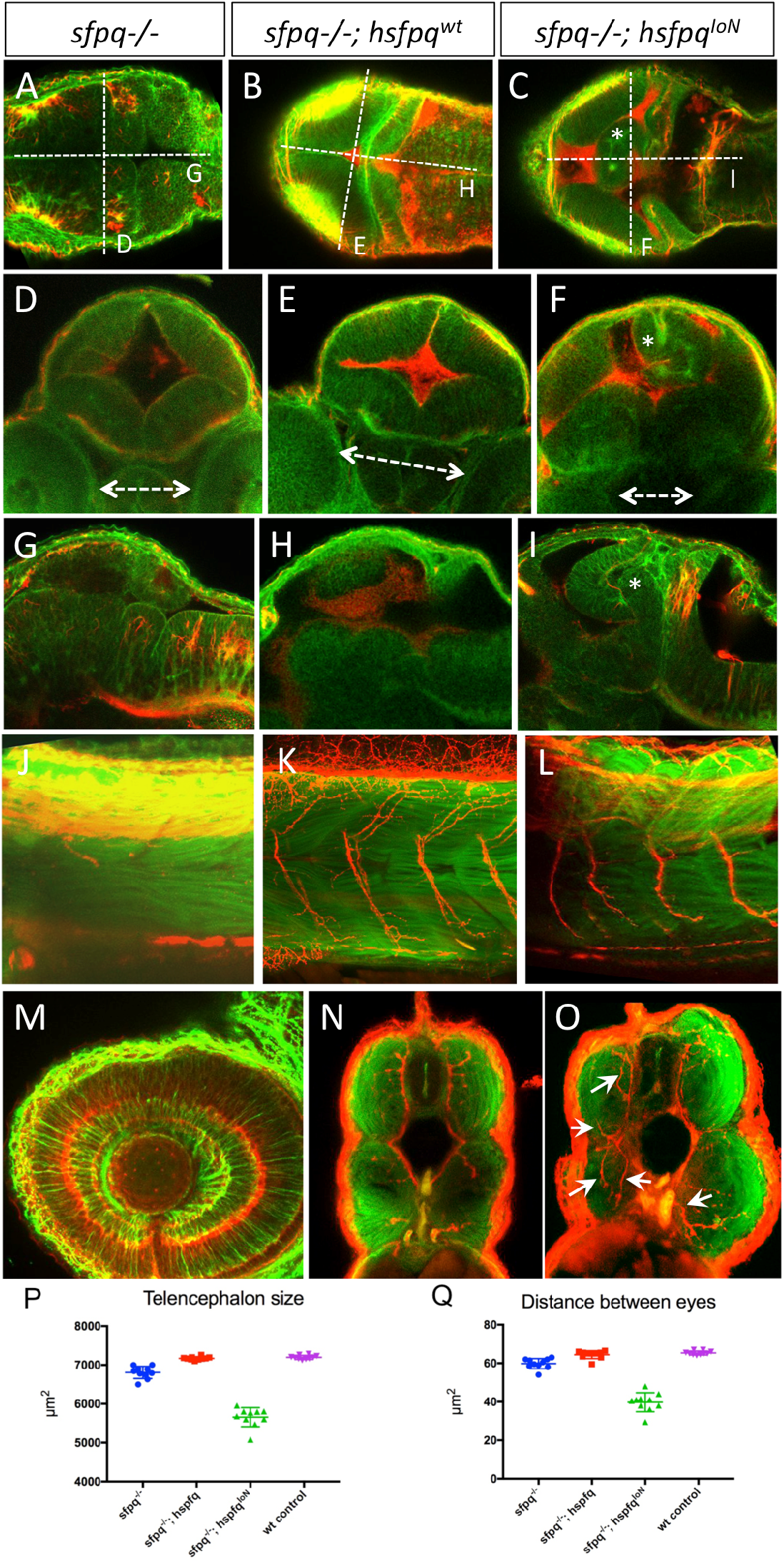
Phenotypic characterisation of *sfpq-/-* zebrafish embryos rescued by ubiquitous expression of human *SFPQ^S660N^* gene. **A-I:** Dorsal (A-C), transverse (D-F, double arrow showing distance between eyes), and lateral (G-I) views of the midbrain/hindbrain Isthmus at 48hpf showing dramatic misfolding (*) in embryos expressing the S660N variant (C, F, I) but not in the mutant rescued by expression of wildtype human SFPQ (B, E, H). **J-L, N, O:** Lateral views (J-L) and transverse sections (N, O) of trunk motor innervation in wildtype (K, N) or S660N variant (L, O) rescued 48hpf embryos. M: Lateral view of the eye in the S660N rescued animal, showing no defect in retinal organisation. **P, Q**: Quantification of telencephalic size and optic distance measured at 48hpf in ten wildtype, (purple), *sfpq-/-* uninjected (blue), *sfpq-/-* injected with wildtype human *SFPQ* (red) or with human *SFPQ* S660N variant (green) embryos, Phalloidin staining in green and acetulated tubulin in red.

**Figure 3.**
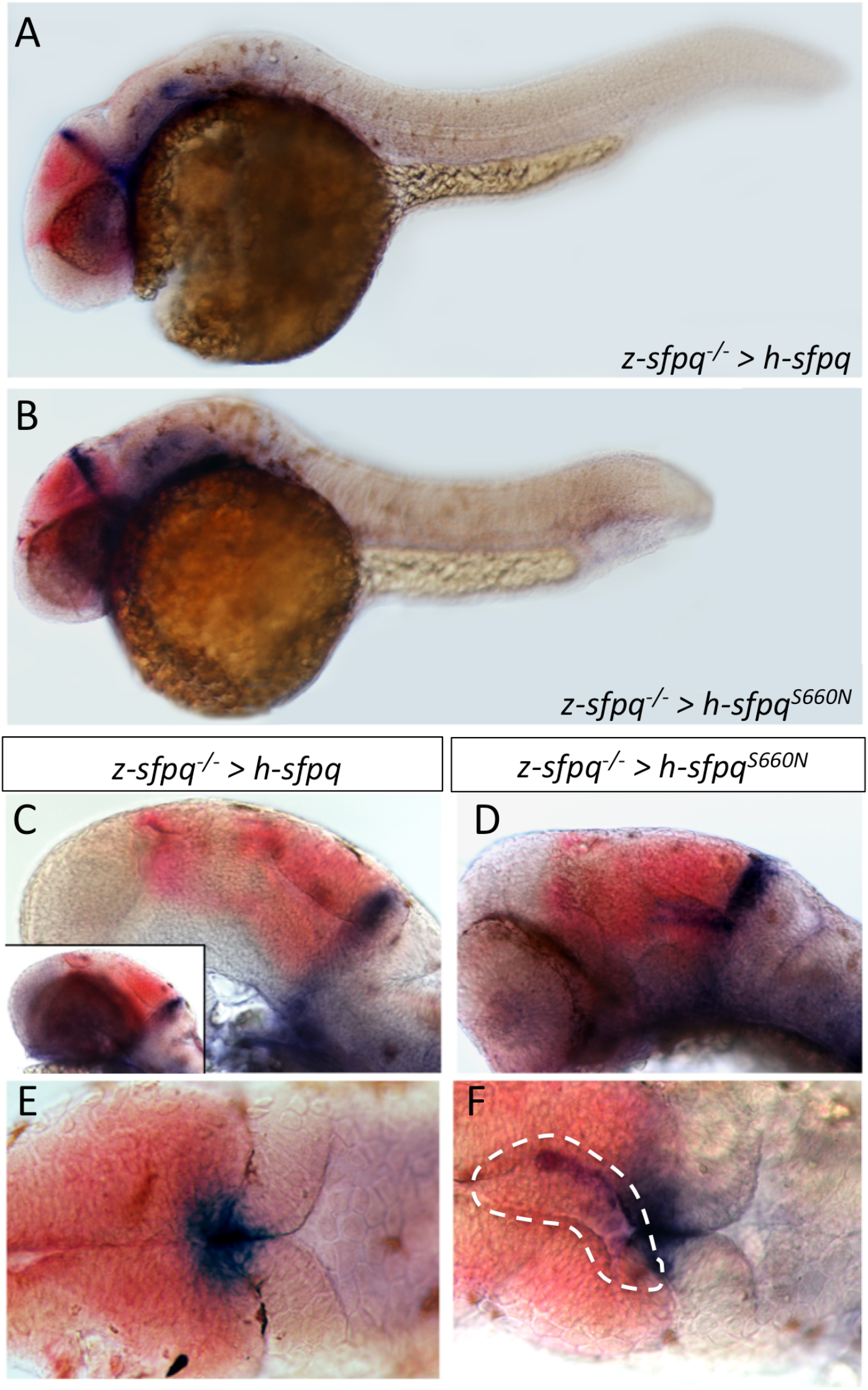
Tissue identity of the ectopic ventricular brain mass. **A-B:** Lateral view of whole sfpq-/-embryos rescued by wildtype (A) or S660N (B) human *SFPQ*, showing *Otx2* (red) and *Gbx2* (blue) expression. Note that the variant-rescued embryos have most often a shorter body. **C-F:** Lateral |(C, D) and dorsal (E, F) close-up view of the midbrain/hindbrain boundary. Inset in C shows the head before eye dissection. This was not possible to achieve in the variant-rescued embryos due to greater adherence of the retina to the rest of the brain. Dotted line in F delineate the ectopic ventricular mass.

All phenotypic analyses of the zebrafish embryos were done without prior knowledge of the patient pathologies. The defects found are strikingly similar to the developmental problems of the patient as suggested by the MRI data.

We then questioned whether the mutated protein was normally localized in the embryonic neurons. We injected gfp-tagged *hsfpq* or *hsfpq^S660N^* RNA into an incross of *sfpq*^+/−^ zebrafish at the one-cell stage and assessed localization using antibody staining at 36 hpf. We found that the hSFPQ^S660N^ protein is very robustly detected in all cell nuclei, similar to wildtype hSFPQ (Figure 4A, 4B). In addition, however, we detected large SFPQ^S660N^ puncta in a proportion of axons imaged (Figure 4A, 4C), substantially brighter and bigger than in the wildtype, indicating an abnormal organization of SFPQ protein complexes in these axons. The puncta were apparent in 4/19 embryos tested, all showing rescued null mutant morphology, indicating that these large *hsfpq^S660N^* puncta only appear in an *sfpq*^−/−^ background. Embryos injected with wildtype hSFPQ did not exhibit visible axonal puncta at the same magnification.

**Figure 4.**
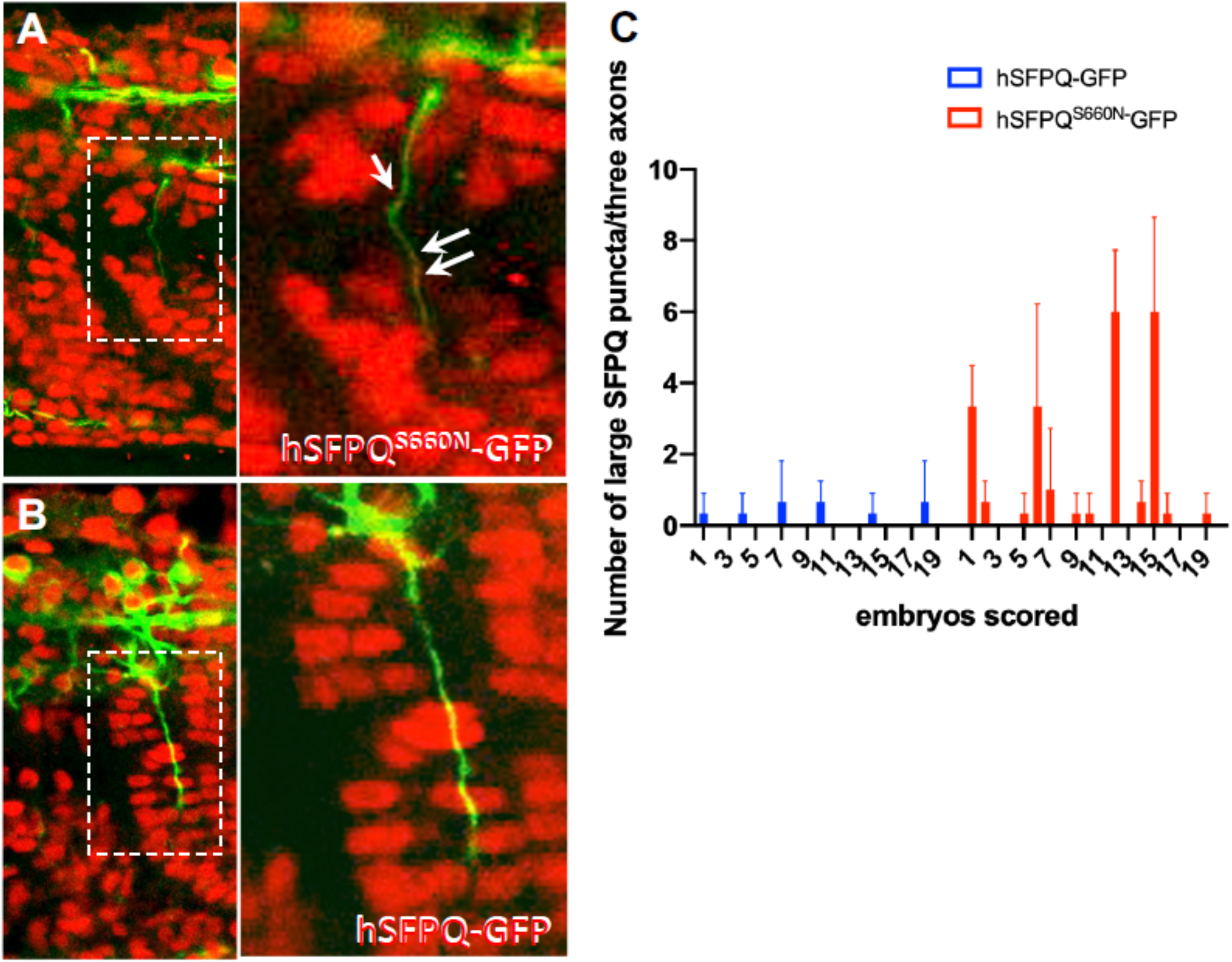
Distribution of human SFPQ protein in rescued zebrafish mutants. **A, B:** Lateral views of confocal imaging of trunk innervation at the level of somite 16 showing human SFPQ expression (red) in 36hpf *sfpq-/-* zebrafish. Axonal tracts stained with acetylated tubulin antibody (green). Arrows showing large puncta of protein in motor axons. **C**: Quantification of large puncta in three axons (somite 15, 16, 17) per embryo in 20 individuals at 36hpf, per rescue condition (wildtype or S660N).

## Conclusions

Taken together, these genetic and functional studies confirmed the vital role of the mRNA splicing factor *SFPQ* for nervous system development and function. They more importantly demonstrate the deleterious effect of the p.Ser660Asn missense mutation, affecting the ability of the SFPQ protein to maintain normal hindbrain and midbrain separation and normal axonal growth and branching during embryonic development. The same function is likely to be conserved across species, thus supporting the conclusion of a set of deleterious neurological consequences of *SFPQ* biallelic p.Ser660Asn mutation in humans. Further animal model work remains to be done to fully understand the molecular dysfunction associated with this variant and how *SFPQ*-related changes in transcription and pre-mRNA splicing regulation may possibly alter downstream brain iron homeostasis and potentially contributing to different degenerative and neurological human phenotypes.

## Acknowledgements

We gratefully acknowledge the family for the enthusiastic collaboration to this study.

---

**Supplementary Figure 1.**
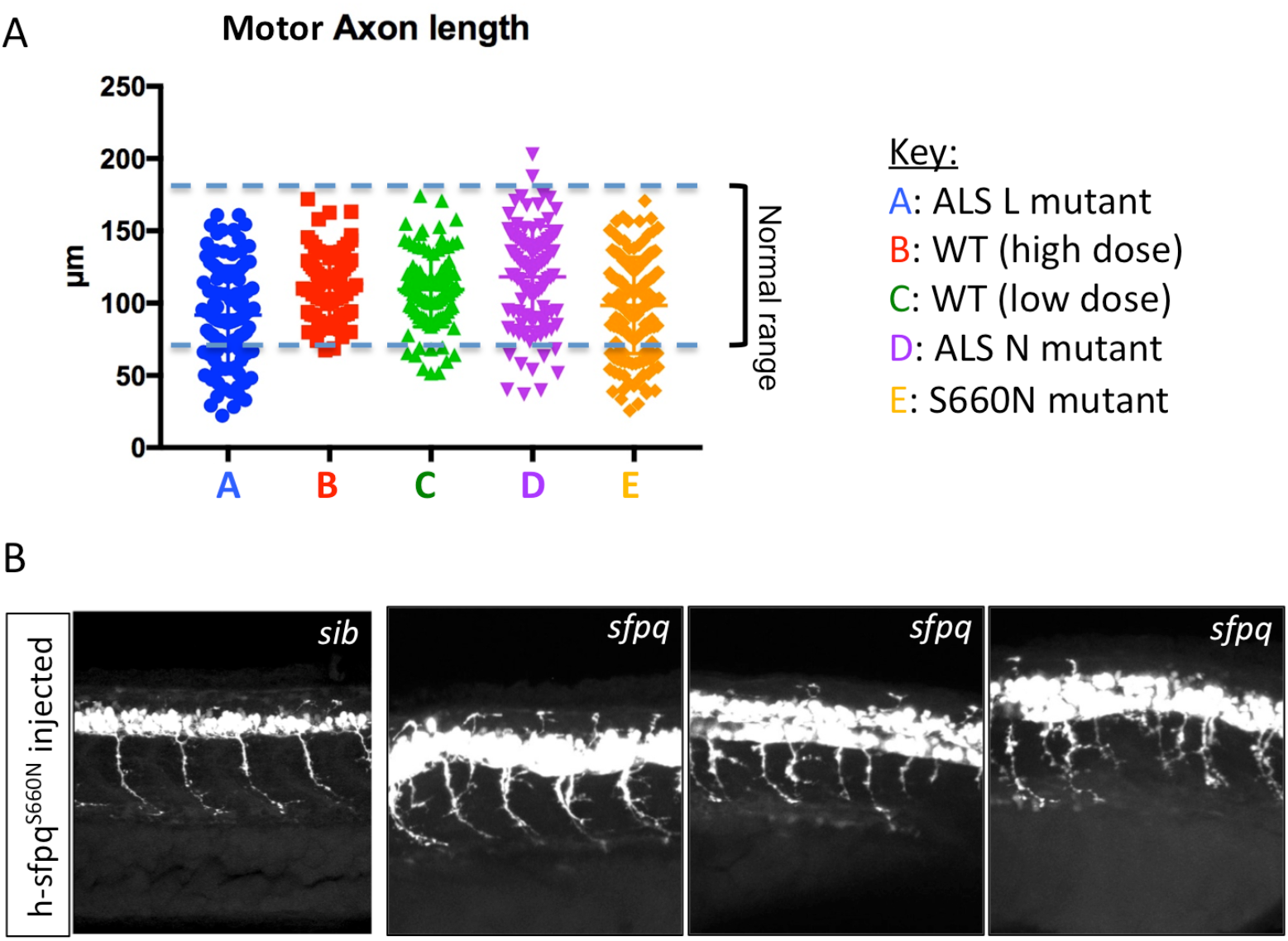
Double-blind phenotypic assessment of zebrafish SFPQ null mutant rescued by various human SFPQ variants. **A:** Length of motor axons in the progeny of *sfpq*^+/−^ cross (1/4 of population is null and 3/4 heterozygous or homozygous wildtype) injected with hsfpq RNA, either wildtype (WT at low dose: 150 pg/embryo or high dose: 250 pg/embryo, to control for phenotype due to overexpression of normal proteins), or with N533H, L534I, or S660N mutations at 150pg/embryo. **B:** Lateral views of spinal cord motor neurons in embryos injected with SFPQ^S660N^ RNA at 48 hpf.

